# Genomic Prediction in Family Bulks Using Different Traits and Cross-Validations in Pine

**DOI:** 10.1101/2021.03.10.434809

**Authors:** Esteban F. Rios, Mario H. M. L. Andrade, Marcio F.R. Resende, Matias Kirst, Marcos D.V. de Resende, Janeo E. de Almeida Filho, Salvador A. Gezan, Patricio Munoz

## Abstract

Genomic prediction (GP) integrates statistical, genomic and computational tools to improve the estimation of breeding values and increase genetic gain. Due to the broad diversity in biology, breeding scheme, propagation method, and unit of selection, no universal GP approach can be applied in all crops. In a genome-wide family prediction (GWFP) approach, the family bulk is the basic unit of selection. We tested GWFP in two loblolly pine (*Pinus taeda* L.) datasets: a breeding population composed of 63 full-sib families (5-20 individuals per family), and a simulated population with the same pedigree structure. In both populations, phenotypic and genomic data was pooled at the family level *in silico*. Marker effects were estimated to compute genomic estimated breeding values at the individual (GEBV) and family (GWFP) levels. Less than six individuals per family produced inaccurate estimates of family phenotypic performance and allele frequency. Tested across different scenarios, GWFP predictive ability was higher than those for GEBV in both populations. Validation sets composed of families with similar phenotypic mean and variance as the training population yielded predictions consistently higher and more accurate than other validation sets. Results revealed potential for applying GWFP in breeding programs whose selection unit are family bulks, and for systems where family can serve as training sets. The GWFP approach is well suited for crops that are routinely genotyped and phenotyped at the plot-level, but it can be extended to other breeding programs. Higher predictive ability obtained with GWFP would motivate the application of GP in these situations.

## Introduction

Genomic (Elshire et al. 2011), statistical (Meuwissen et al., 2001; Gianola et al., 2009), and computational advances have allowed significant increases in genetic gain by applying genomic prediction (GP) in breeding programs across several species (e.g., Hayes et al., 2009; Fe et al., 2015, 2016; Gezan et al., 2017; de Bem Oliveira et al., 2020; Amadeu et al., 2020). Taking advantage of the ever-reducing cost of molecular markers (Wetterstrand, 2020), the concept of GP was derived (Meuwissen et al., 2001) as an alternative method to marker-assisted selection (MAS). Genomic prediction utilizes a dense panel of molecular markers covering the whole genome to predict genomic estimated breeding values (GEBV) of individuals with no phenotypic records (Meuwissen et al., 2001). Traditional GP pipelines involve developing a training set (TS), for which available genotypic and phenotypic data is fitted to build a prediction model. This model is later used to predict GEBV of selection candidates in a validation set (VS), composed of individuals that are genotyped but not phenotyped. Cross-validation schemes are implemented taking sub-samples from the TS to calibrate the model and then use the model into the remaining part of the TS to estimate and evaluate its predictive ability, i.e. the correlation between GEBVs and phenotypic values (Perez-Cabal et al., 2012).

Genomic prediction has been quickly adopted in animal breeding (Hayes et al., 2009) due to readily accessible genomic data, large reference populations with accurate pedigree records, and the impossibility of phenotyping sex-linked traits (Stock and Reents, 2013). In dairy cattle, GP can double the genetic gain compared to selection based on progeny test (Xu et al., 2020). On the contrary, the application of GP in plants has been lagging behind due to less accessible high-throughput genotyping methods, lack of accurate pedigree records, and the wide range of variation in life cycle, ploidy level, and mating systems found in plants (Hough et al., 2013). All these plant-specific characteristics are key factors affecting predictive ability in GP due to their influence in breeding methods, effective population size, population structure, and linkage disequilibrium (Lin et al., 2014). Pioneer studies implementing GP in plants were performed in mayor crop species with traditional hybrid selection such as maize (Massman et al., 2013; Combs and Bernardo, 2013) and trees (Resende et al., 2012; Kumar et al., 2012), or variety selection in self-pollinating species (Poland et al., 2012). Genomic prediction showed to be a powerful tool to achieve higher genetic gain in plant breeding in many other species (Crossa et al., 2017; Lara et al., 2019; de Bem Oliveira et al., 2020; Esfandyari et al, 2020). Large commercial breeding companies have been applying GP; however, the success of the process depends strongly on the species architecture and the breeding program scheme (Xu et al., 2020; Voss-Fels et al., 2019)

Several species are bred as populations of large full or half-sib families, and commercially used as populations of different levels of relationship (i.e. synthetic cultivars) as in some forage species, such alfalfa (*Medicago sativa* L.) (Annichiarico et al., 2015; Biazzi et al. 2017) and ryegrass (*Lolium perenne* L.) (Fe et al., 2016; Cericola et al., 2018). In these species, the family (full or half-sibs) is the basic unit for phenotyping (e.g. plot-level measurement for yield rather than plant level) and selection. Thus, due to the mating system nature (allogamy), individual plants are of limited interest because commercial varieties represent a homogenous population composed of heterozygous individuals (Poehlman, 1987). Also, it is not straightforward to link phenotypic data collected on individual spaced-plants to plot-based swards in crops such as forage and turfgrass, which are mostly allogamous (Poehlman, 1987), and single-plant performance has been shown to poorly predict plot-based data (Wang et al., 2016). Therefore, the application of genome-wide family prediction (GWFP) would be advantageous for traits that are phenotyped using family pools in swards or plots. The phenotypic data collection at the plot level could be extended to other organisms grown and evaluated in families, such as turfgrasses (*Lolium perenne* L.), forages (*Medicago sativa* L.), sugarcane (*Saccharum officinarum* L.), cassava (*Manihot esculenta* L.), and to aquaculture species such as shrimp (*Litopenaeus vannamei*) (Barbosa et al., 2012; Torres et al., 2019; Pembleton et al., 2018, Jia et al., 2018, Wang et al., 2017). The application of GWFP has already been reported for crops that are bred and farmed as family pools, such as cross-pollinated forage species (Fe et al., 2015, 2016; Guo et al., 2018; Cericola et al., 2018, Annichiarico et al., 2015; Biazzi et al. 2017; Jia et al., 2018).

The GWFP approach considers family-pools as the measurement unit. Here, both allele frequencies and phenotypic records are expressed as a single average record of a given family. Therefore, the additive genetic variance in full-sib families is half of the additive variance between individuals (i.e. only 50% of the genetic variation is exploited in GWFP), which would result in higher predictive ability when compared to GEBV (Ashraf et al. 2014). Despite the initial efforts to test the predictive ability of GWFP using empirical data, there is a need to explore further implementation of GWFP in breeding schemes. As a first aspect, it is essential to compare the predictive ability of GEBV vs. GWFP models, and to develop strategies to combine both approaches. For this, datasets that contain family structures but genotyped and phenotyped at the single plant level are ideal. Another aspect is the understanding of the influence that family/pool size and phenotypic variances in training/validation sets have in the predictive ability for various traits.

In order to evaluate these aspects, two loblolly pine (*Pinus taeda* L.) populations were studied: a) an observed breeding population composed of 63 families (CLONES_real), and b) a simulated population that reproduced the same pedigree as CLONES_real. The objectives of this study are: i) to identify the minimum number of individuals per family required to calculate allele frequency and phenotypic mean values with reasonable accuracy; ii) to investigate the effect of contrasting phenotypic mean and variance between training and validation sets on predictive ability; and iii) to assess the predictive ability of GEBV and GWFP. Loblolly pine is not normally bred in family pools, but existing real and simulated datasets were used to compare GEVB and GWFP approaches.

## Materials and Methods

### Observed population

The loblolly pine (*Pinus taeda* L.) population known as “comparing clonal lines on experimental sites” (CCLONES_real) has previously been used for predicting performance of individual trees (Resende et al., 2012). In this study, GWFP was tested by pooling individual trees belonging to the same full-sib family. The population is composed of 923 individuals from 70 full-sib families obtained by crossing 32 parents in a circular diallel mating design with additional off-diagonal crosses (Baltunis et al., 2007). The number of individuals per family ranged from 1 to 20, with an average of 13 trees per family (standard deviation = 5). In this study, families with less than five individuals were removed, and 63 full-sib families were used for analyses. Data collection was described in detail in Resende et al. (2012) and Munoz et al. (2014). In summary, all 923 genotypes from CCLONES_real was phenotypically characterized in three replicated studies and was genotyped using an Illumina Infinium assay (Illumina, San Diego, CA; Eckert et al. 2010) with 7,216 SNPs, each representing a unique pine EST contig. In the current study, four traits representing growth, quality, and diseases were selected based on their narrow-sense heritability and genetic architecture as reported by Resende et al. (2012). These correspond to: a) lignin concentration (Lignin) (*h*^*2*^ = 0.11, polygenic trait), b) tree stiffness (Stiffness) at year 4 (km^2^/sec^2^) (*h*^*2*^ = 0.37, polygenic trait), c) rust susceptibility (Rust) caused by *Cronartium quercuum* Berk. Miyable ex Shirai f. sp. *Fusiforme* (*h*^*2*^ = 0.21, oligogenic trait), and d) diameter at breast height (Diameter) at year six (cm) (*h*^*2*^ = 0.31, polygenic trait).

### Simulated Population

A simulated population (CCLONES_sim) exhibiting similar genetic properties as CCLONES_real was also considered in this study. Genomic prediction approaches using individual trees were previously explored using this synthetic population (de Almeida Filho et al., 2016, 2019). For its simulation, the base population was created (G0 = 1,000 diploid individuals) by randomly sampling 2,000 haplotypes from a population with an effective size of *N*_*e*_ = 10,000 and a mutation rate of 2.5 × 10^−8^. Then, the 10% highest phenotypic values from G0 were selected and randomly mated to generate the first breeding generation (G1). From G1, 42 individuals were selected and used in a circular diallel mating design that reproduced the pedigree as in CCLONES_real (G2), comprised of 923 individuals and 71 full-sib families. However, only 63 families, with more than five individuals, were used in this study. Subsequently, 42 individuals were selected from G2 and used in crosses to the next generation (G3, CCLONES_sim_prog), a population composed of 1,176 individuals and 71 families. Only the 63 families with more than five individuals were used for analyses. The simulated genome had 12 chromosomes and 5,000 polymorphic loci, and only the scenario exhibiting an absence of dominance (*d*^*2*^ = 0.0) and *h*^*2*^ = 0.25 were used for analyses in this study. Two traits with different genetic architectures were simulated: i) oligogenic: 30 QTL were sampled from a gamma distribution with rate 1.66 and shape 0.4, with positive or negative QTL effects (Meuwissen et al., 2001), and ii) polygenic: 1,000 QTL were used, and their additive effects were sampled from a standard normal distribution (Hickey and Gorjanc, 2012).

### Phenotypic and Genotypic Data Pooling

In both populations, phenotypic and genotypic data were pooled at the family level *in silico*. The phenotypic data were averaged across all individuals belonging to the same full-sib family; therefore, the average phenotypic value by family was used as the response for all analyses. In the case of the genomic data, the allele frequency (p) was calculated for each SNP per family, considering the reference allele (A) as follows:

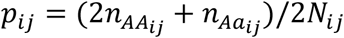

Where *p*_*ij*_ refers to the allele frequency for SNP *i* in the *j* family; *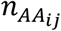* and 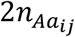 are number of individuals with genotype AA and Aa respectively for SNP *i* in the family *j*; *N*_*ij*_ are number of individuals in family *j* with non-missing genotype data for SNP *i*. Missing values for allele frequency were imputed at the family level using the average allele frequency for that given SNP across families. Markers were excluded from analyses when more than 50% of the families exhibited missing values, and SNPs were not removed based on minor allele frequency. A total of 4,740 polymorphic SNPs (CCLONES_real) and an average of 5,000 polymorphic SNPs for CCLONES_sim and CCLONES_sim_prog (average across simulated replicates) were used in the analyses.

### Number of Individuals per Family

A total of 10 families from CCLONES_real with at least 15 individuals were selected to evaluate the minimum number of individuals required to estimate allele frequency and phenotypic family means with the most reasonable accuracy. Families were specifically selected to represent segregation ratios (1:1 and 1:2:1) for 10 SNPs. Allele frequencies per family and family phenotypic means were calculated varying the number of individuals per family from one to 15. These values were used to compute the squared deviations between the mean value obtained with *i* number of individuals (*i* = 1 to 15) and the mean value obtained with the entire family (15 individuals), under the assumption that 15 individuals per family provide accurate estimates of allele frequencies and phenotypic mean in our families. This assumption can be validated using the concept of genetic representativeness, given by the effective population size (Ne). The estimator of the Ne within a full sib family is given by Ne = [2n/(n+1)] (Resende and Barbosa, 2006). The maximum (when n goes to infinite) Ne within a full sib family is 2. With n equal to 15 individuals the Ne is 1.88, which is 94% of this maximum of 2.

### Statistical Methods

Marker effects were estimated at the individual (GEBV) and family (GWFP) levels with two distinct whole-genome regression approaches using the package BGLR (Perez and de los Campos, 2014) in R (R Development Core Team, 2018): i) *Bayes B* which considers that markers have heterogeneous variances, i.e., many loci with no genetic variance and a few loci explain a large portion of the genetic variation (Meuwissen et al., 2001; Perez and de los Campos, 2014); and ii) *Bayes RR* a Bayesian method that assumes common variance across all loci; therefore, SNPs with the same allele frequency explain the same proportion of variance and have the same shrinkage effect (Gianola, 2013; Perez and de los Campos, 2014).

In total, 20,000 Markov chain Monte Carlo iterations were used, of which the first 5,000 were discarded as burn-in, and every third sample was kept for parameter estimation. We fitted the following model for individual and family models:

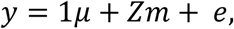

Where ***y*** is the vector of the averaged phenotype by family in the case of GWFP and by individual in the multiple clones in the case of GEBV, *μ* is the overall mean fitted as a fixed effect, ***m*** is the vector of random marker effects, and ***e*** is the vector of random error effects, **1** is a vector of ones, and ***Z*** is the incidence matrix indicating allele frequencies in the case of GWFP (ranging from 0 to 1), and marker dosage (0, 1 and 2) for GEBV.

After fitting the model described above for each trait, the GEBV and GWFP of family/individual *j* (g_j_) were obtained using the following expression:

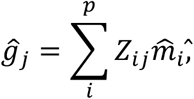

Where *i* is the allele frequency/marker dosage of the *i-*th marker on family/individual *j*, and *p* is the total number of markers, and 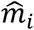, is the estimated effect of *i*-th SNP.

### Creating Training/Validation Sets Using Contrasting Phenotypes

Phenotypic values for each trait in both populations were sorted and divided into three classes: the smallest 10%, the largest 10%, and values between both extremes. Five validation sets were created for each trait using these phenotypic classes: a) Low: 10% families with the lowest phenotypic values; b) High: 10% families having the highest values; c) Low+High: combining four families from Low and three families from High; d) Middle: seven families showing phenotypes around the population mean, e) Combined: two families from Low, two families from High, and three families from Middle. For the populations Low+High (c), Middle (d), and Combined (e), three replicates were created by taking random samples from each phenotypic class. The other 56 families were used as training sets to build prediction models.

### Split-Families as Training/Validation Sets

All families with more than ten individuals (59 in total) were randomly split into two equivalent size groups. One group of individuals phenotypic and genotypic data were pooled at the family level and used as the training set (TST) for GWFP models. The other group of individuals was used as the validation population (VST) based on two approaches: i) predicting the performance of individuals trees not included in the TST (GWFP_Fam_Ind), and ii) pooling individuals at the family level to predict performance of families composed of individuals not included in the TST (GWFP_Fam_Fam).

### Prediction in the Following Generation in CCLONES_sim

The GP models were developed by using the G2 CCLONES_sim population as the TST. These training models were used and validated in the G3 generation using individuals (GEBV) and family pools (GWFP), and models were assessed by calculating predicted ability and prediction accuracy. Predicted ability was estimated by calculating a Pearson’s correlation between the phenotypic values and the estimated breeding values, and prediction accuracy was estimated by calculating a Pearson’s correlation between the real breeding value and the estimated breeding value.

### Model Validation and Predictive Ability

Prediction models for GEBV and GWFP were validated using 10-fold cross-validation and leave-one-out (LOO) approaches. For the 10-fold CV, data was randomly partitioned into ten subsets, and TST populations were created with 90% of the families/individuals, while the remaining 10% of families/individuals were used as VST. This scheme was repeated until the ten subsets were used as VST. In the LOO approach, models were constructed using *N*_*T*_ -1 families (where *N*_*T*_ = is the total number of families) in the TST. The validation set was the single family not included in the training group. This scheme was repeated *N*_*T*_ times until all 63 families were used as the TST.

Each time the models were fitted using a different VST, the model’s predictive ability was estimated calculating a Pearson’s correlation between the observed/simulated phenotypes and the GWFP/GEBV estimates for the families/individuals included in the VST.

### Data Availability

All phenotypic and genotypic data utilized in this study have been previously published as a standard data set for development of genomic prediction methods (Resende et al. 2012; de Almeida Filho et al., 2016). Simulated data available from the Dryad Digital Repository: http://dx.doi.org/10.5061/dry-ad.3126v.

## Results

### Number of Individuals per Family

The minimum number of individuals per family was calculated assessing allele frequency and phenotypic mean deviations using families with at least 15 individuals. For genotypic and phenotypic data, the lowest number of individuals needed to accurately estimate allele frequency and family means was six (Figure 1). Allele frequency deviations (Figure 1 A-D) and mean phenotypic deviations (Figure 1 E-F) indicated that families with less than six individuals were not providing accurate estimates of the family’s genotypic and phenotypic means in both populations. We assumed that the observed values based on 15 individuals per family provides with a reasonable estimation of allele frequency and phenotypic mean for a diploid species. Therefore, all 63 families with six or more individuals were used for further analyses in this study. Both populations showed similar trends for the genotypic and phenotypic estimates (Figure 1). The average allele frequency deviations were lower for SNPs exhibiting a 1:1 ratio in both populations (Figure 1 A and C), compared to SNPs segregating into a 1:2:1 ratio (Figure 1 B and D). For phenotypic data, CCLONES_sim showed slightly smaller deviations, especially for a lower number of individuals (Figure 1 F), compared to CCLONES_real for the trait diameter (Figure 1 E). Other traits in CCLONES_real exhibited a similar behavior (data not shown).

**Figure 1.**
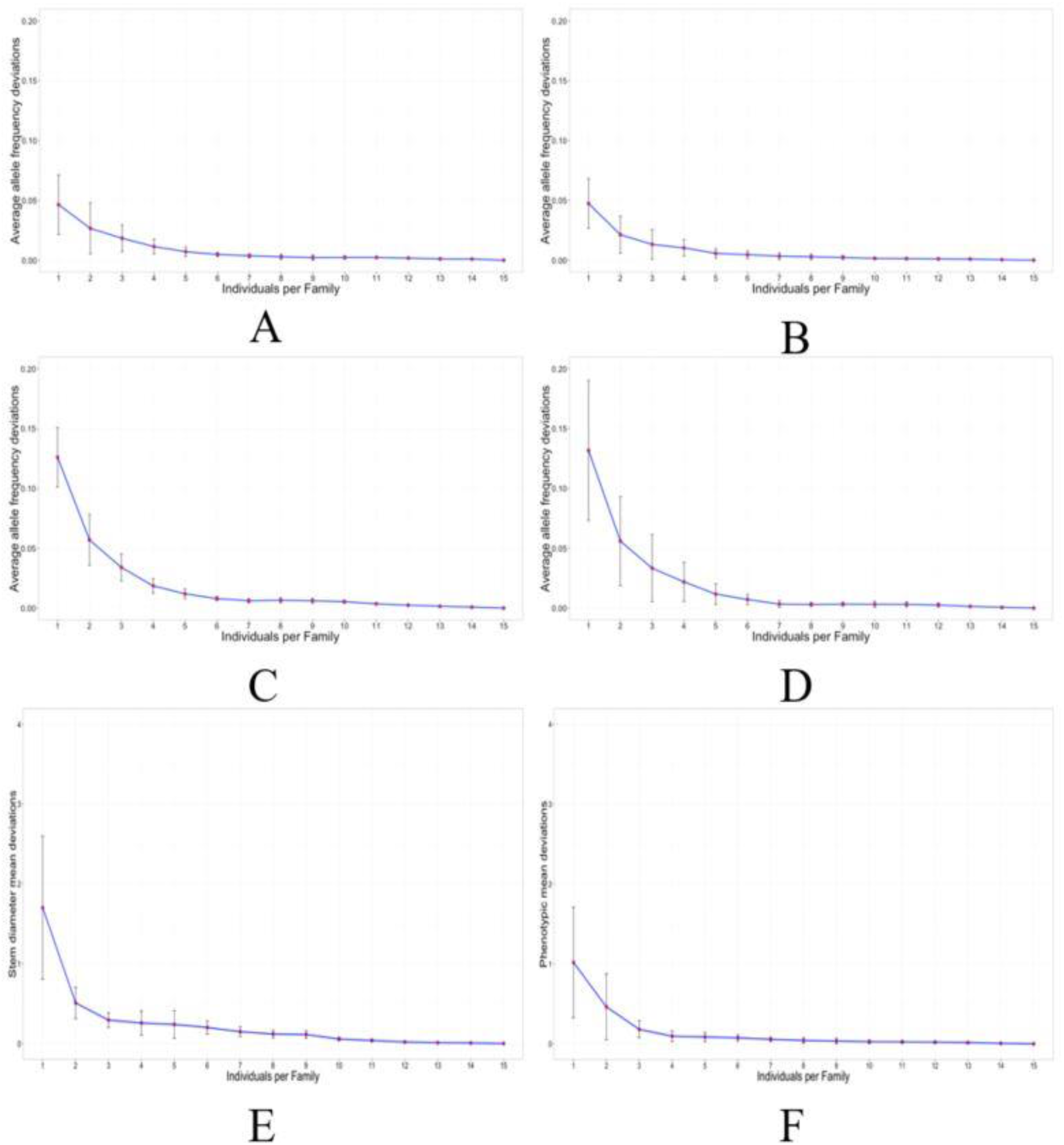
Average allele frequency deviation (A-D) and family mean phenotypic deviation (E-F) in CCLONES_real (A, C and E) and CCLONES_sim (B, D and F) calculated by increasing the number of individuals from 1 to 15. Five families exhibiting genotypic segregation ratios 1:1 (A and B) and 1:2:1 (C and D) for single nucleotide polymorphisms were included in the analysis. The CCLONES-real phenotypic deviation is for the trait stem diameter (E).

### Statistical Method and Cross-Validation

Two Bayesian statistical methods (*Bayes B* and *Bayes RR*) and two cross-validation approaches were used to test the predictive ability of GWFP in four traits measured in CCLONES_real (Figure 2). Both statistical methods yielded high and similar predictive abilities for the four traits (Figure 2 A and B). However, standard errors for predictive ability were larger with the LOO approach (Figure 2 A and B). Additionally, GWFP predictive abilities obtained with the LOO approach were slightly lower than for the 10-fold cross-validation scheme (except for trait Stiffness) (Figure 2 A and B). Therefore, the 10-fold cross validation approach was selected to perform further analyses.

**Figure 2.**
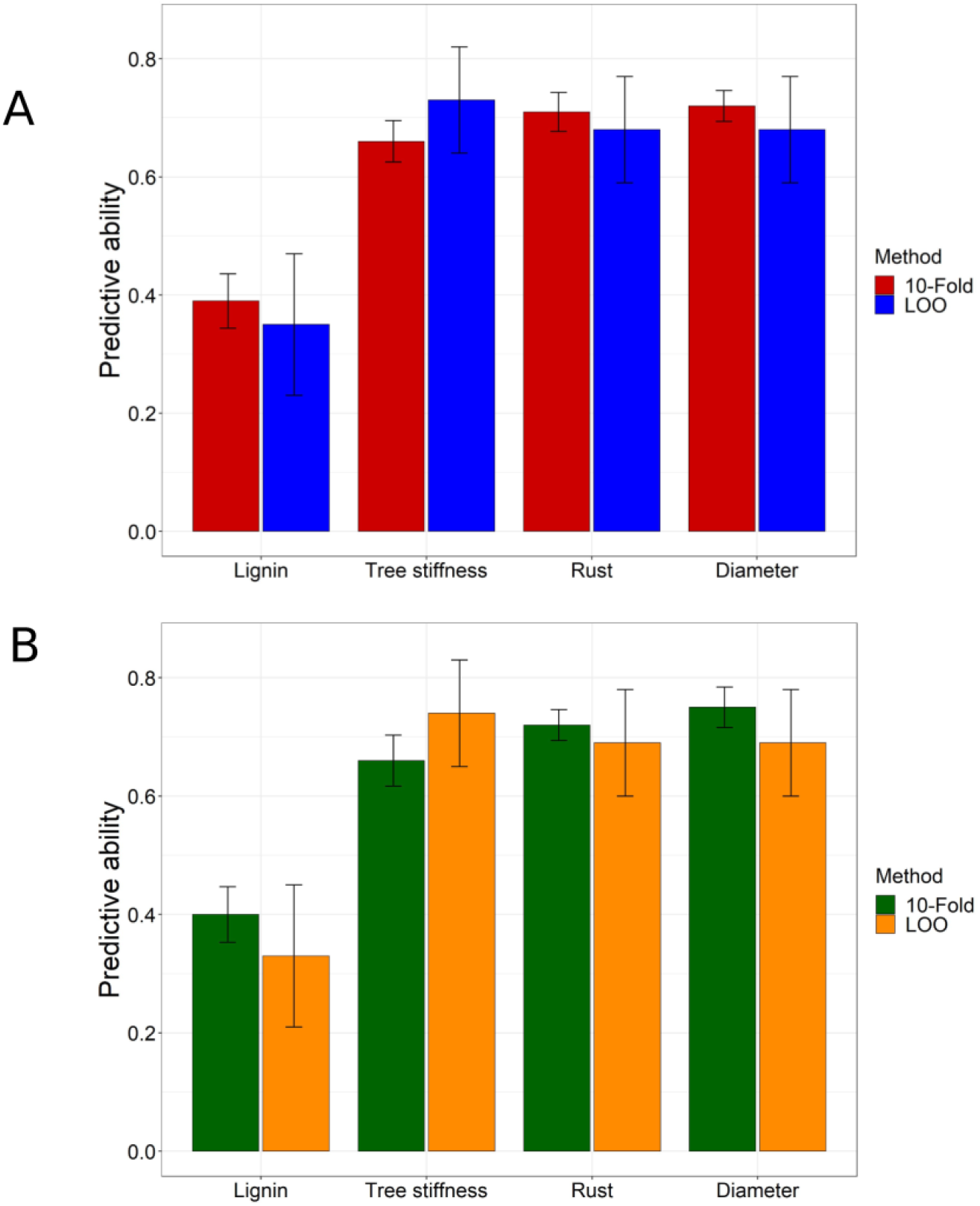
Average predictive ability using family pools (GWFP) in four traits in the loblolly pine breeding population CCLONES obtained with 10-fold and leave-one-out (LOO) cross validation schemes using *Bayes B* (A) and *Bayes RR* (B).

### Predictive Ability of GWFP Using Training/Validation Sets with Contrasting Phenotypes

The effect of phenotypic data in the predictive ability of GWFP was explored by creating five VST’s using contrasting sets of phenotypic data between TST and VST (Figure 3 A). The predictive ability for GWFP for all traits were least accurate and had larger standard errors when the VST was composed of families exhibiting small and large phenotypic values (bottom and top classes) (Figure 3 B). When VST’s were composed of families exhibiting phenotypes corresponding to the middle class, predictive ability increased for all traits, but standard errors were still large (Figure 3 B). As expected, there was an increase in predictive ability and a large reduction in standard errors when VST’s were composed of families showing similar phenotypic mean and variance to the TST, corresponding to the classes “Low+High” and “Combined” (Figure 3 B).

**Figure 3.**
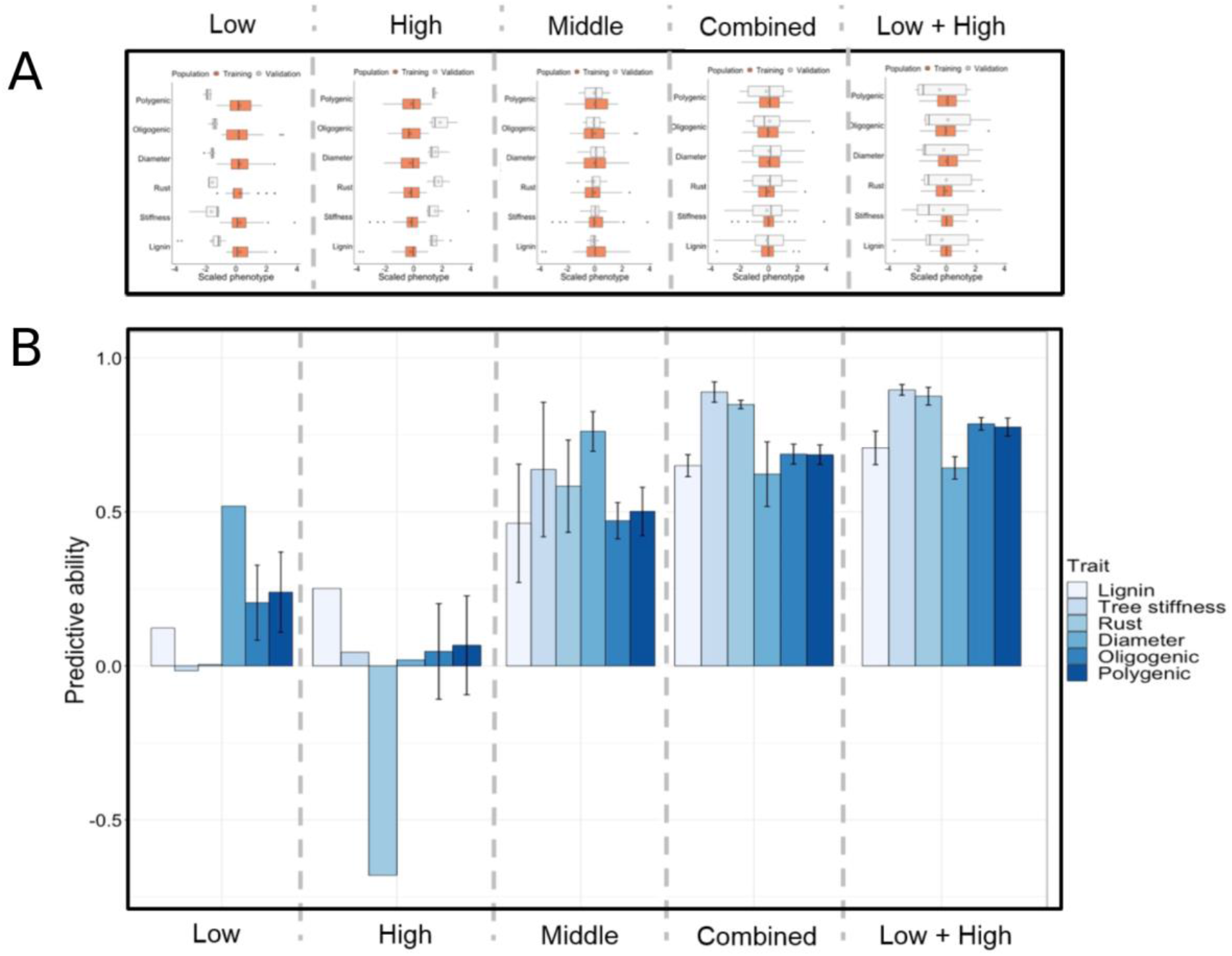
Phenotypic distribution for testing (orange) and validation (white) sets for fours traits measured the CCLONES_real population and two traits simulated using CCLONES_sim (A). Average predictive ability obtained with *Bayes B* using genome wide family prediction (GWFP) for four traits in the CCLONES_real (lignin, stiffness, rust and diameter), and two traits with different genetic architecture (Oligogenic and Polygenic) in the CCLONES_sim populations (B). Five scenarios were tested by creating training (56 families) and validation (7 families) populations using phenotypic data: i) **Low**: validation set is composed of 7 families with lowest phenotypic records; ii) **High**: validation set is composed of 7 families with highest phenotypic records; iii) **Middle**: validation set is composed of 7 families with phenotypic records similar to the family mean; iv) **Combined**: 2 families from Low, 2 families from High and 3 families from Middle; and v) **Low + High**: 4 families from Low and 3 families from High.

### Predictive Ability of GEBV and GWFP

Predictive ability obtained with *Bayes B* using different methods and schemes (Table 1) is presented in Figure 3 for the 63 families from both populations. The traditional GP approach with individuals in the TST and VST (GEBV) was contrasted with predictive ability obtained with the family-based (GWFP) method following a 10-fold cross validation scheme. The scenarios GWFP_Fam_Ind and GWFP_Fam_Fam were run only once because CCLONES (real and simulated) had a limited number of individuals per family (Figure 4).

**Table 1.** Scenarios implemented to design training and validation sets to test predictive ability of genomic prediction models.

| Scenario | Set |  |
| --- | --- | --- |
|  | Training | Validation |
| GEBV | 830 individuals | 93 individuals |
| GWFP | 56 families | 7 families |
| GWFP_Fam_Ind | 59 families | 422 individuals |
| GWFP_Fam_Fam | 59 families | 59 families |
| GWFP_Low | 56 families | 7 families with lowest phenotypic values |
| GWFP_High | 56 families | 7 families with highest phenotypic values |
| GWFP_Low_High | 56 families | 7 families, 4 lowest and 3 highest phenotypic values |
| GWFP_Middle | 56 families | 7 families with values similar to the overall mean |
| GWFP_Combined | 56 families | 7 families (2 Low, 2 High and 3 from Middle scenarios) |

GEBV: genomic estimated breeding value.
GWFP: genome-wide family prediction.
CV: cross validation.

**Figure 4.**
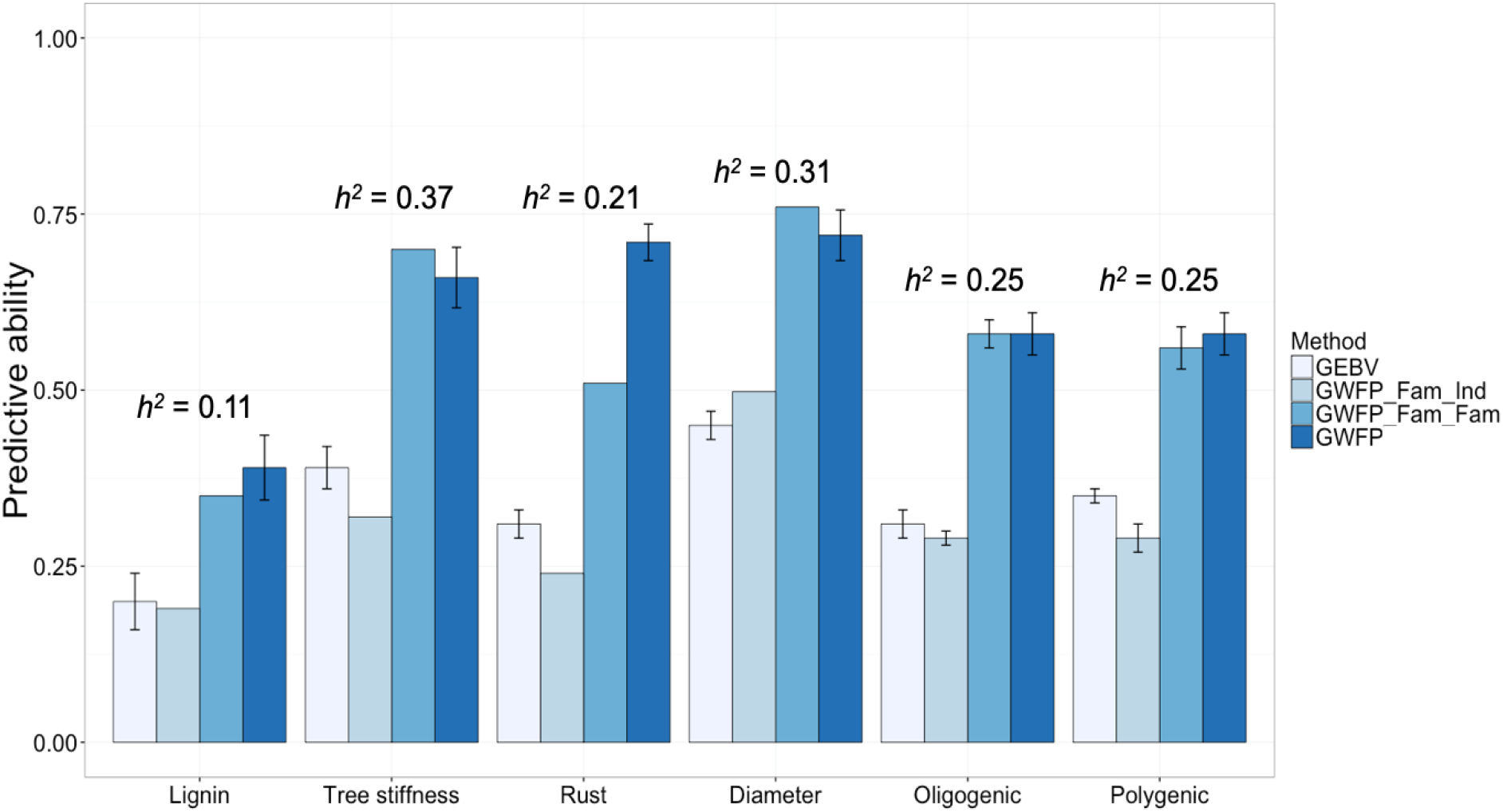
Average predictive ability obtained with *Bayes B* for four traits in CCLONES-real (lignin, tree stiffness, rust and stem diameter), and two traits with different genetic architecture (Oligogenic and Polygenic) in the CCLONES_sim populations using different genomic prediction methods. GEBV: genomic estimated breeding values individual trees; GWFP_Fam_Ind: genome-wide family prediction using 59 family pools as training set, while different individuals from the same families were used as validation set; GWFP_Fam_Fam: genome-wide family prediction using 59 family pools as the training and validation population, but different full-sib individuals were pooled in both sets; GWFP: genome-wide family prediction using 63 family pools in a 10-fold cross validation scheme. Narrow-sense heritability (*h*^*2*^) estimated at the individual level (Resende et al., 2012).

Predictive ability was always greater for GWFP methods in both populations and all traits, except for the scenario GWFP_Fam_Ind that showed similar or lower accuracy than GEBV for most traits (Figure 4). Additionally, predictive ability was greater for traits with higher heritability (Figure 4). Specifically, GWFP provided predictive abilities at least 40% greater than traditional GEBV for most of the traits in both populations. Moreover, GWFP_Fam_Fam exhibited similar or greater predictive ability than GWFP for most traits in both populations, except for rust (Figure 4). Both sets of traits from the simulated CCLONES population exhibited very similar accuracies for all schemes (Figure 4).

### Predictive Ability and accuracy of GEBV and GWFP in the Following Generation

Accuracy and predictive ability of GEBV and GWFP were obtained with the prediction models built with the CCLONES_sim (G2) population as the TST, and models were validated in the following generation (G3). The GEBV showed higher accuracy than GWFP for the oligogenic trait, and similar accuracy for the polygenic trait (Figure 5). Predictive ability for the oligogenic and polygenic traits were higher for GWFP (Figure 5). Additionally, greater predictive ability and accuracy were observed for the oligogenic trait, and the difference between accuracy and predictive ability was greater for the oligogenic trait (Figure 5).

**Figure 5.**
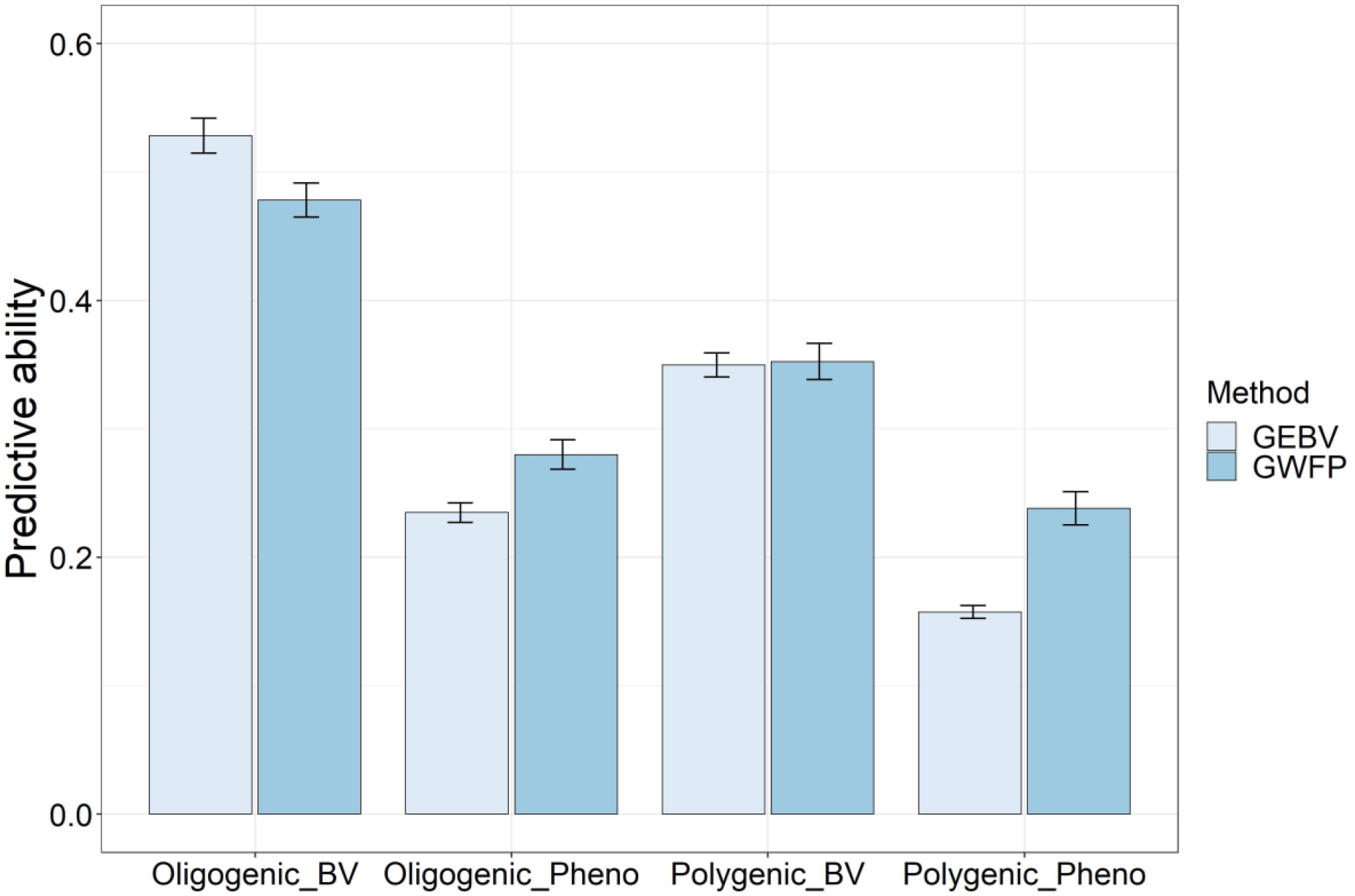
Average predictive ability and accuracy obtained with *Bayes B* for two traits with different genetic architecture (Oligogenic and Polygenic) in the CCLONES_sim_progeny population, obtained with individual (GEVB) and family-pooled (GWFP) genomic prediction methods. Predictive ability calculated as the correlation between estimated breeding and phenotypic values are denoted as _Pheno, and accuracy as the correlation between estimated and true breeding values as _BV.

## Discussion

We quantified the predictive ability of GWFP in real and simulated loblolly pine breeding populations for different traits and cross-validation approaches. Moderate to low predictive ability values were obtained with the traditional GP approach, as previously reported for both populations, using individual trees as the basic phenotypic and genotypic unit (Resende et al., 2012; de Almeida Filho et al., 2016). In general, GWFP outperformed GEBV in the predictive ability for most traits; including the predictive ability for the oligogenic and polygenic traits in CCLONES_sim when using the following generation (G3) as the VST.

### Family Size

The size and structure of the training population affects the accuracy of GP models (Van Raden et al., 2009; Daetwyler et al., 2010; Habier et al., 2010; Grattaglia and Resende, 2011; Edwards et al., 2019; de Bem Oliveira et al., 2020). In our study, the size of the TP refers to the number of families and the number of individuals within a family. The number of families was fixed and limited to 70 families, so we did not focus on studying the effect of a variable number of families. However, the minimum number of individuals per family to obtain reasonable accurate estimates of family allele frequency and family phenotypic mean was found to be six. When studying the effect of size and composition of training population in blueberry (*Vaccinium* spp.), De Bem Oliveira et al. (2020) found a high predictive ability using six individuals per family for some traits. Thus, in their study family variance was accurately represented with six individuals per family in this autotetraploid species. Using the estimator of the *Ne* within a full sib family, given by *Ne* = [2n/(n+1)] (Resende and Barbosa, 2006), the maximum (when n goes to infinite) *Ne* within a full sib family is 2. With n equal to 6 individuals the *Ne* is 1.71, which is 86% of the maximum 2. So, n = 6 appears adequate to represent genetically a full-sib family, corroborating our results.

The effect of number of individuals within families on accuracy of GP models was also demonstrated in perennial ryegrass (Pemblenton et al., 2016; 2018). The authors stated that 48 to 60 individuals per population are necessary to accurately represent the genetic diversity within a ryegrass population. As an allogamous species, multiple parents are used to create synthetic populations in perennial ryegrass, hence multiple individuals with a high number of loci in heterozygosis are contributing to the variation in the synthetic population. Perennial ryegrass is commonly bred using families and GWPF has been exploited in the species for various traits (Fe et al., 2015, 2016; Guo et al., 2018; Cericola et al., 2018).

Simulation studies with variable numbers of families and individuals per family would help identify the optimum training population sizes for GWFP. Generally, a larger training population (more families in the training population) yield higher accuracy (Voss-Fels et al., 2019; de Bem Oliveira et al., 2020), but this is associated with higher costs. Therefore, the definition of the optimum number of families, and number of individuals per family are a crucial point for the genomic prediction process. Fe et al. (2015) studied the effect of the number of families in the accuracy of genomic prediction for heading date in ryegrass; the authors found high accuracies with a low number of families (<100). The authors showed that increasing the number of families to 500 leads to higher accuracy, and more than 500 families did not yield to significant improvement.

### Statistical Methods and Cross-Validation Scheme

Models considering different Bayesian methods were similar in predicting GEBV in traits measured in the real breeding population and the simulated population in this study. Resende et al. (2012), reported a slightly greater predictive ability in the real population for rust incidence with Bayesian methods over RR-BLUP, because fewer genes with large effects control this trait. De Almeida Filho et al. (2016), using the simulated population, reported a slightly lower predictive ability in the oligogenic trait using *Bayes RR* than *Bayes B*. In the present study, *Bayes B* and *Bayes RR* were tested to compare their performance in GWFP because prior distributions and assumptions for both methods are contrasting (Perez and de los Campos, 2014). Our results showed that both *Bayesian* methodologies were very similar in predicting family-pools, even for rust incidence in the real population and for the oligogenic trait in the simulated population.

Both cross-validation schemes, LOO and 10-fold, produced similar results in predicting GWFP with a slight advantage for the 10-fold scheme, due to the large variation in the LOO scheme. Resende et al. (2012) reported similar results with the real data set for GEBV, wherein 10-fold and LOO resulted in no significant differences in their predictive ability. Also, similar predictive abilities between the 10-fold and LOO scheme have been reported in wheat (*Triticum aestivum* L.) (Edwards et al., 2019).

### Predictive Ability of GWFP Using Contrasting Phenotypes

When the families in the VST had phenotypic values outside the range of phenotypes presented in the TST (bottom and top classes), lower and much more variable predictive abilities were obtained. Interestingly, higher predictive abilities were obtained when families in the VST had the same phenotypic range as the TST. The impact of the phenotypic variance on prediction was demonstrated by Edwards et al. (2019), which reported that the accuracy of genomic prediction in wheat showed higher predictions for crosses (validation set) with higher phenotypic variance. Würschum et al. (2017) reported equivalent results in triticale (x *Triticosecale* Wittmack), in which higher accuracy was detected for the traits of plant height and biomass in cases in which families with a large phenotypic variation were included in the training/validation set population.

The differences in predictive ability among the scenarios for phenotypic values in the VST could also be related to the composition of the TST’s. For the extreme scenarios (Low and High), the TST’s did not have the extreme phenotypic values and alleles frequencies, which could have resulted in poor estimations of markers effects. Studying the optimization process for genomic prediction in wheat, Norman et al. (2018) showed that the genomic prediction accuracy could be improved, in cases when TST and VST are not related, by increasing the genetic diversity in the TST.

### Predictive Ability of GEBV and GWFP

Predictive ability was always greater for GWFP methods than GEBV in both the real and simulated populations and for all traits, except when the model was built with family pools, and individual performance was predicted (GWFP_Fam_Ind) (Figure 4). Although the full sib families average explores only half of additive genetic variance, the error variance is mitigated with larger number of observations due progeny replication, when compared with single observations (Hallauer et al. 2010). Then, this higher precision of phenotypic value in family bulks could explain the higher accuracy in genomic prediction of families.

The higher accuracy in the GWFP method was expected since the additive genetic variance explored in this method is just 50% of the additive genetic variance compared to the GEBV, which leads to a higher accuracy and heritability (Casler et al. 2008; Ashraf et al. 2014). Besides, relatedness between the TST and the VST also influence the predictive ability. The relationship between the TST and VST has a crucial role in the model predictive ability (Lorenz & Smith 2015; de Bem Oliveira et al., 2020), it can help explain the higher predictive ability found in the GWFP_Fam_Fam and GWFP, compared to the GEBV and GWFP_Fam_Ind.

Nevertheless, the predictive ability for most traits obtained with GWFP_Fam_Ind scheme was of the same order of magnitude compared to GEBV, except for the traits stiffness and rust. Therefore, using the numbers from this study as example, considering the significant reduction in costs incurred in DNA extraction and genotyping 56 families (TST for GWFP), instead of 844 individuals (TST for GEBV), the approach GWFP_Fam_Ind could still be an affordable option for implementing GP in breeding programs that select individual plants, but have limited budgets to phenotype and genotype all individuals in the training set.

Reduced investments to implementation of genomic prediction with higher predictive ability accuracies can be obtained with the GWFP approach compared with GEBV. A larger number of families can be included in the models, which, for the present population, would likely result in higher predictive abilities as reported in perennial ryegrass for heading date (Fe et al., 2015). Additionally, including more than 10 individuals per family will reduce the sampling variability of the allele frequency and phenotypic mean, resulting in higher genomic accuracies (de Bem Oliveira et al. 2020).

### Application of GWFP in a breeding program

Breeding cycles can take several years in perennial crops, and phenotyping costs could be high for critical production and quality traits. Genomic prediction has the power to shorten the time of a breeding process, which leads to a higher genetic gain per unit time, and can allow a reduction in phenotyping process and costs (Grattaglia and Resende, 2011; Crossa et al., 2017; Voss-Fels et al., 2019). Genotyping cost has been decreasing, allowing the extensive use of molecular markers in breeding programs. However, in some cases, breeders need to genotype a large number of individuals (>10,000) to implement GP in their programs, increasing costs significantly (Voss-Fels et al., 2019). The high genotyping costs due to large population sizes can make it impracticable to implement GP in minor crops, particularly in public breeding programs.

For breeding programs with limited budgets, the GWFP can be an alternative to GEBV due to the reduction in phenotypic and genotypic costs to develop prediction models. GWFP has been used in several forage species that are bred in family bulks and whose phenotyping for critical traits is conducted at the sward/plot level (Fe et al., 2015, 2016; Guo et al., 2018; Cericola et al., 2018, Annichiarico et al., 2015; Biazzi et al. 2017; Jia et al., 2018). In a GEBV approach, the data (phenotypic and genotypic) is collected at the individual level and models are built to estimate the performance of individuals (Figure 6-A) (Resende et al., 2012; de Almeida Filho et al., 2016, 2019). The GEBV requires significant more resources (labor, economic, computational) to collect and analyze data. Under a GWFP approach, the number of genotypic samples (bulked DNA and a single sequencing effort per family) will be the exact number of families, representing a significant reduction in the number of samples compared to the traditional GEBV process (Fig. 6-B). The phenotyping process will also be performed at the family/plot level, which is the ideal scenario for critical traits in some crops such as forage and turfgrass species.

**Figure 6.**
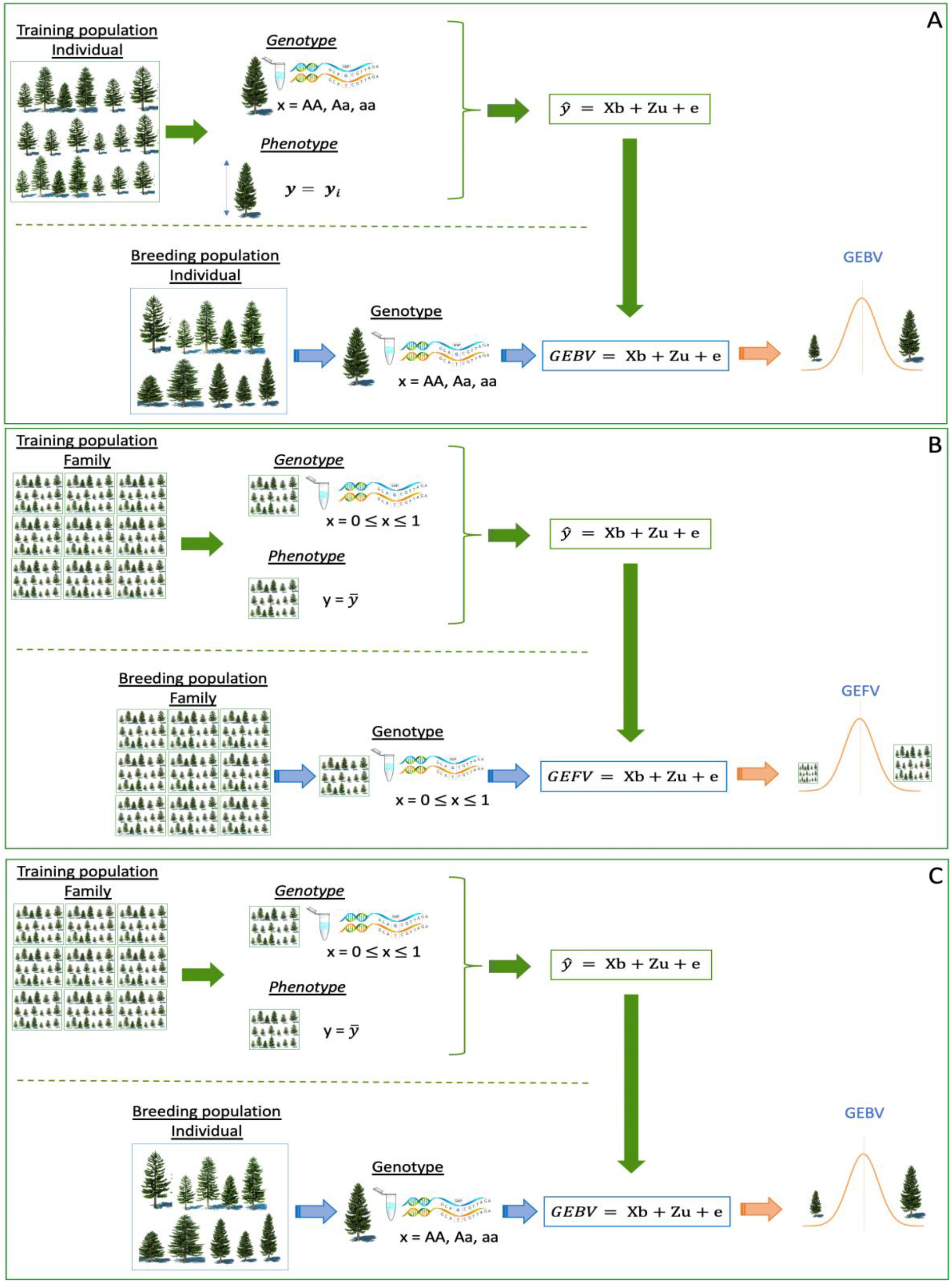
Scheme for the different genomic prediction scenarios: A - GEBV: genomic estimated breeding values for individual trees; B – GWFP_Fam_Fam: genome-wide family prediction for families prediction; C – GWFP_Fam_Ind: genome-wide family prediction applied in the selection of individuals.

Breeders may also be interested in employing the GWFP_Fam_Ind approach, where family bulks are used as training set, but individuals are the selection unit (Figure 6-C). In this study, the GWFP_Fam_Ind approach showed similar accuracy to GEBV for most traits, with the addition of lower needs for phenotypic and genotypic data for the model development. Finally, GWFP models could be exploited in scenarios when remnant seeds might be available for the same family, and the goal would be to predict the performance of the family or individuals within the family. The remaining seeds from the selected families can be used later to test their merits in further replicated field trials. For perennial allogamous crops, families used in the TST set can be used as a new crossing block to start a new selection cycle.

## Conclusion

Despite the limitation in number of families and number of individuals per family tested in this study, less than six individuals per family produced inaccurate estimates of family phenotypic performance and allele frequency. Validation sets with similar phenotypic mean and variance as the TST set showed greater predictive ability and more accurate predictions consistently across traits. These results revealed great potential for using GWFP in breeding programs that select family bulks as the selection unit, GWFP is well suited for crops that are routinely genotyped and phenotyped at the plot-level. The GWFP approach can also be extended to breeding schemes where family bulks can serve as training sets, while individuals are the selection target.

